# Deep mutational scanning of the plasminogen activator inhibitor-1 functional landscape

**DOI:** 10.1101/2021.04.15.440003

**Authors:** Zachary M. Huttinger, Laura M. Haynes, Andrew Yee, Colin A. Kretz, David R. Siemieniak, Daniel A. Lawrence, David Ginsburg

## Abstract

The serine protease inhibitor (SERPIN) plasminogen activator inhibitor-1 (PAI-1) is a key regulator of the fibrinolytic system, inhibiting the serine proteases tissue- and urokinase-type plasminogen activator (tPA and uPA, respectively). Missense variants may render PAI-1 non-functional through misfolding, leading to its turnover as a protease substrate, or to a more rapid transition to the latent/inactive state. Deep mutational scanning was performed to evaluate the impact of amino acid sequence variation on PAI-1 inhibition of uPA using an M13 filamentous phage display system. The effects of single amino acid substitutions on PAI-1’s functional inhibition of its canonical target proteases, tPA and uPA, have been determined for only a small fraction of potential mutations. To construct a more comprehensive dataset, a mutagenized PAI-1 library, encompassing ∼70% of potential single amino acid substitutions, was displayed on M13 filamentous phage. From this library, the relative effects of 27% of all possible missense variants on PAI-1 inhibition of urokinase-type plasminogen activator were determined using high-throughput DNA sequencing with 826 missense variants demonstrating conserved inhibitory activity and 1137 resulting in loss of PAI-1 function. Comparison of these deep mutational scanning results to predictions from PolyPhen-2 and SIFT demonstrate the limitations of these algorithms, consistent with similar reports for other proteins. Comparison to common human PAI-1 gene variants present in the gnomAD database is consistent with evolutionary selection against loss of PAI-1 function. These findings provide insight into structure-function relationships for PAI-1 and other members of the SERPIN superfamily.

## INTRODUCTION

Plasminogen activator inhibitor-1 (PAI-1, *SERPINE1*) is a member of the <u>se</u>rine <u>pr</u>otease <u>in</u>hibitor (SERPIN) protein superfamily (1). SERPINSs function as irreversible inhibitors that covalently bind the active site of their target proteases (2). All inhibitory SERPINs share a common mechanism wherein a flexible reactive center loop (RCL) extends outside the central structure of the molecule serving as a cleavable peptide “bait” for the target protease (Fig. 1A) (3). The SERPIN and protease form an acyl intermediate when the active site serine acts as a donor nucleophile to bind to the carbonyl carbon of the P1 residue within the RCL(4,5). However, the RCL rapidly inserts into central β-sheet A before the hydrolysis reaction is completed, carrying the tethered protease to the opposite side of the molecule in a pole-to-pole transition (6). This structural transformation stabilizes the protease-SERPIN complex and renders both the SERPIN and the serine protease no longer active (2,7,8). If the rate of RCL insertion is slower than the resolution of the acyl intermediate, complete proteolysis of the scissile bond occurs, with the SERPIN then functioning as a substrate instead of an inhibitor (9). Either of these mechanisms is possible when a SERPIN interacts with a given protease. The specificity of a SERPIN for its target protease is in part driven by the probability that the inhibitory pathway dominates over the substrate pathway (10). This balance presents a unique challenge when engineering SERPIN variants that inhibit non-canonical target proteases (11,12).

**Figure 1.**
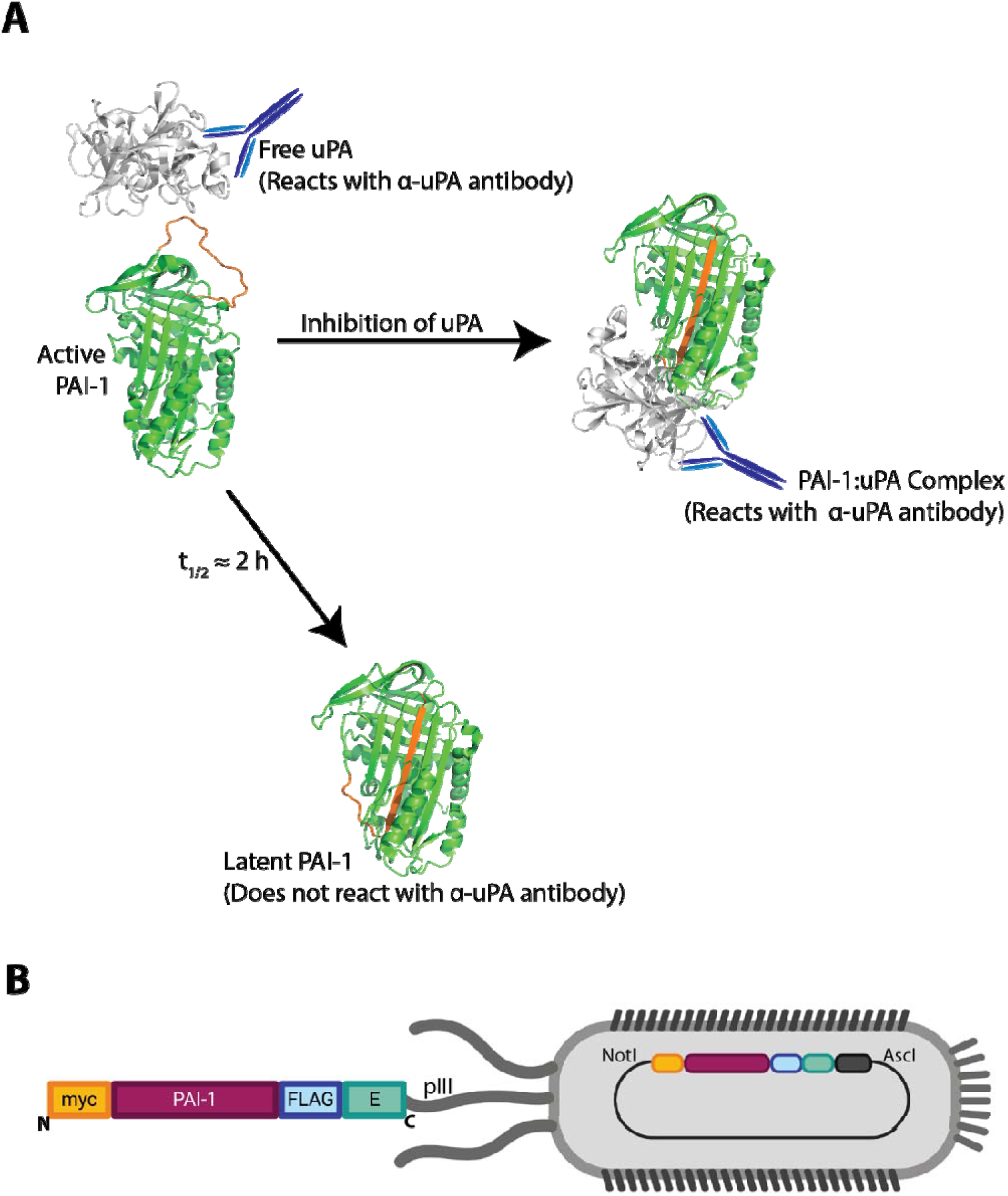
PAI-1 displayed as a fusion protein on the surface of filamentous phage is an inhibitor of uPA. *(A)* Active PAI-1 (PDB: 3PB1) reacts with free uPA (PDB:1W0Z) to form a covalent uPA:PAI-1 complex (54,55). Both free and complexed uPA can be immunoprecipitated using an anti-uPA antibody. Alternatively, PAI-1 can spontaneously relax from its active, metastable state to a low-energy yet chemically inert latent conformation (PDB: 1DVN) (56). PAI-1’s reactive center loop is highlighted in orange. *(B)* Schematic of phPAI-1 displayed as a fusion to the pIII coat protein of M13 filamentous phage, with the N-terminal myc- and C-terminal FLAG-and E-tags highlighted. Created with Biorender.com.

PAI-1 is the principal inhibitor of the serine proteases tissue-type and urokinase-type plasminogen activators (tPA and uPA, respectively)(13,14). tPA and uPA activate the zymogen plasminogen to plasmin (15), the enzyme responsible for the proteolysis of fibrin—the structural backbone of blood clots (16). Human deficiency of PAI-1 results in excessive plasmin generation and a mild to moderate bleeding diathesis (17-21). Likewise, PAI-1 deficient mice are viable with a mild increase in fibrinolytic activity (22). In addition to its critical role in regulating hemostasis, PAI-1has been implicated in a number of other processes, including a link to longevity (23). PAI-1 lacks native cysteine residues and remains functional in the absence of glycosylation, facilitating large-scale production of functionally active, recombinant PAI-1 in bacterial systems (24,25).

The present work couples phage display with high-throughput DNA sequencing (HTS), to measure the effects of multiple missense mutations on PAI-1’s uPA inhibitory function in a massively parallel fashion.

## RESULTS

### Characterization of PAI-1 fusion protein

Selection of a 9:1 mixture of phage displayed PAI-1 (phPAI-1) and VWF A3 domain (phVWF-A3) with uPA (Fig. 2) resulted in at least a five-fold enrichment in phPAI-1 relative to phVWF-A3, indicating that the immunoprecipitation is specific for uPA and uPA:PAI-1 complexes.

**Figure 2.**
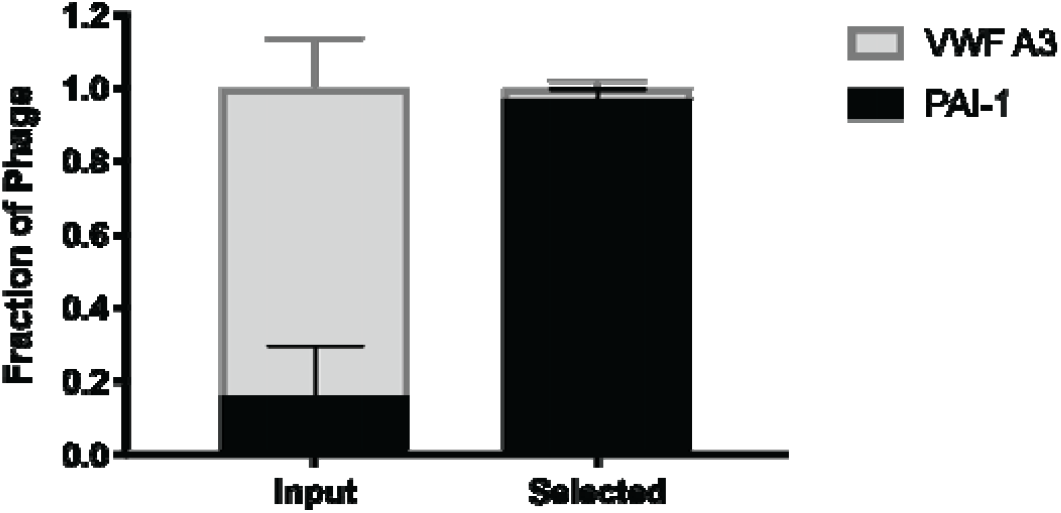
Immunoprecipitation is specific for phPAI-1:uPA complexes. Nine parts phVWF-A3 and one part phPAI-1 (9:1) were combined (*input*, n = 3), incubated with uPA (1.7 nM) for 30 min at 37°C, and selected by immunoprecipitation with an anti-uPA antibody (*selected*, n =3). For each replica, 24 colonies were genotyped by PCR using primers common to both the phVWF-A3 (677 bp) and phPAI-1 inserts (1271 bp) followed by analysis on a 1% agarose gel.

Given the rigorous washing of the immunoprecipitated complex, this enhancement further suggests that PAI-1 expressed as a pIII fusion protein on the phage surface retains its inhibitory activity.

### Characterizatio006E of the mutant library

The phPAI-1 mutant library exhibits a depth of 8.04×10^6^ independent clones with an average of 4.2 ± 1.8 amino acid substitutions/molecule as determined by Sanger sequencing of 13 randomly selected phage clones. HTS demonstrated that 5,117 of the possible 7,201 missense variants (71%) are present in the mutant phPAI-1 library, along with 269 of the 379 possible nonsense mutations (71%). The frequency of DNA sequencing reads for individual amino acid substitutions within the starting library ranged over >10^4^-fold. Additionally, biased amino acid subsitutions were also observed with reduced representation of Met and Trp substitutions (both encoded by only a single codon), compared to Arg, Leu, and Ser substitutions (each encoded by six codons) (Fig. 3).

**Figure 3.**
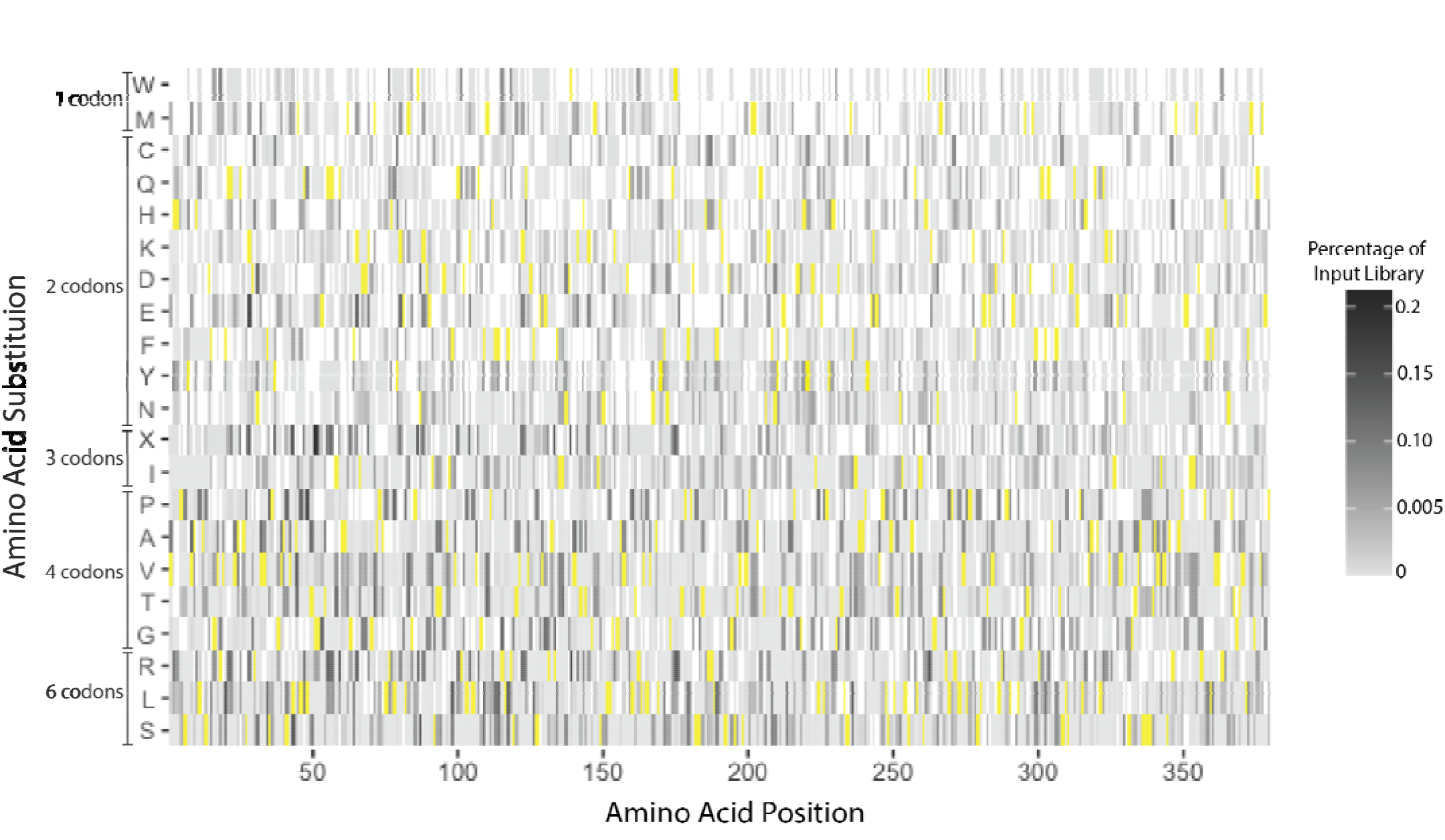
phPAI-1 mutant library generated by error prone PCR includes more than two-thirds of all possible missense mutations. The mutational library contains 71% of all possible missense and nonsense mutations with 27% of all missense variants present with sufficient depth (base mean score > 10, p_adj_ < 0.05) to accurately determine the effects of the mutation on PAI-1 function. The primary amino acid position within PAI-1 is indicated along the x-axis and single amino acid substitutions are listed along the y-axis. WT amino acid residues are indicated in yellow, while missense and nonsense (X) mutations not present in the input library are shown in white. Variants present within the library are shown in grey (see scale) as a percentage of the input library represented by that variant.

To limit the proportion of phPAI-1 variants transitioning to the non-reactive latent conformation, all reactions with uPA were performed immediately following phage production. To limit false positives within the dataset, only those variants with a basemean score (the average of the counts in the input and selected libraries) greater than 10 and an adjusted p-value (p_adj_) < 0.05 were included in further analyses (Fig. 4) (26). Based on these criteria, 1,963 (38%) of the 5,117 missense variants present in the starting library could be scored for uPA reactivity or lack thereof.

**Figure 4.**
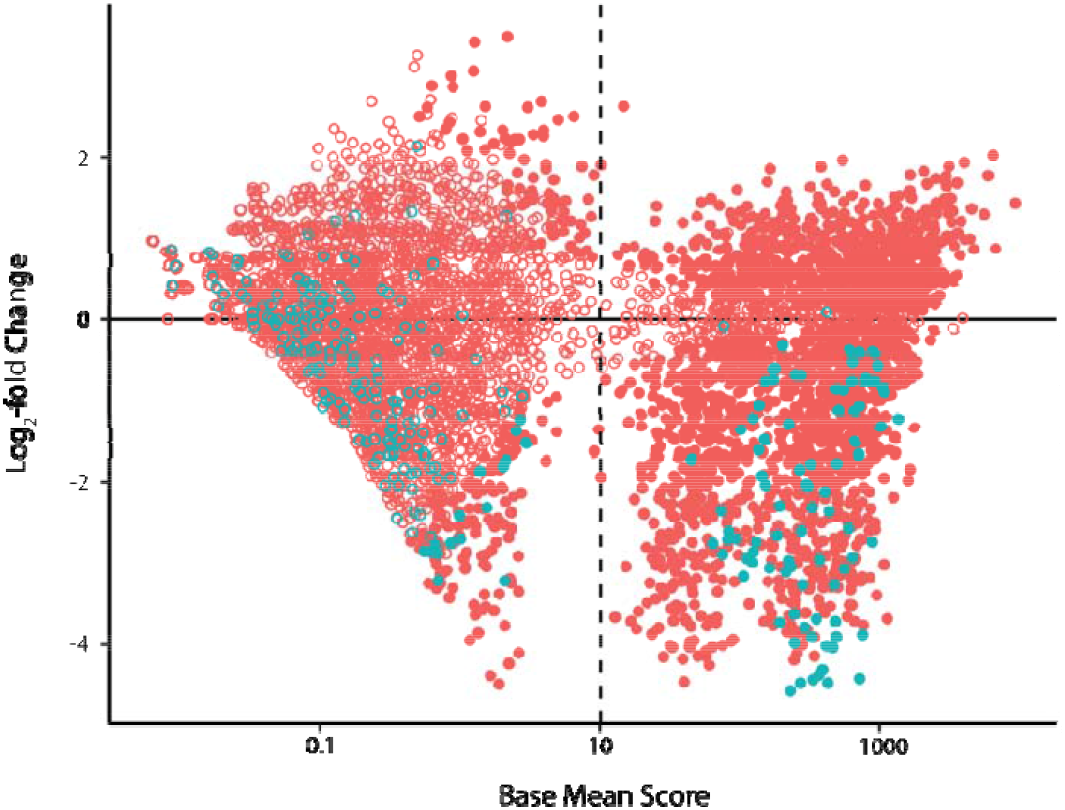
phPAI-1 uPA-selected libraries are enriched in variants that retain their inhibitory function and depleted of those without inhibitory activity. An MA plot (26) of base mean score (average of counts in the input and selected libraries) vs. log_2_-fold change is shown, with missense mutations in pink and nonsense variants in blue. Variants with p_adj_ < 0.05 are shown as closed circles, while those that do not meet this significance threshold are shown as open circles. A base mean score greater than 10 was also set as a threshold for determining significance.

### Massively parallel assessment of variant impact on uPA inhibitory function

Following selection with uPA, 826 PAI-1 missense variants retained the ability to form a complex with uPA, with a range of enrichment scores likely representing varying degrees of inhibitory activity towards uPA (Fig. 5). Similarly, depleted variants (log_2_-fold enrichment score ≤ 0, n = 1137) are broadly classified here as loss-of-function, likely include variants that retain a low level of inhibitory activity towards uPA—again, reflecting that this approach enables the mapping of functional variability with respect to both gain and loss of function (Fig. 5). While WT PAI-1 was enriched 2-fold in the selected versus the input library, other variants were enriched or depleted up to 6- or 23-fold respectively. For comparison, consider two mutations at Ile^91^, I91L and I91N, each representing approximately 0.0005% of the of the input library (Fig. 3). Following selection with uPA, I91L was enriched three-fold, consistent with previous reports (27). In contrast, I91N was depleted three-fold—demonstrating that while the I91L mutation is well tolerated, I91N results in loss of function with respect to uPA inhibition. Of note, the selection method employed here (complex formation with uPA) does not distinguish between the three potential mechanisms for loss-of-function: PAI-1 misfolding, accelerated transition to the inactive latent state, and/or serving as a substrate for uPA.

**Figure 5.**
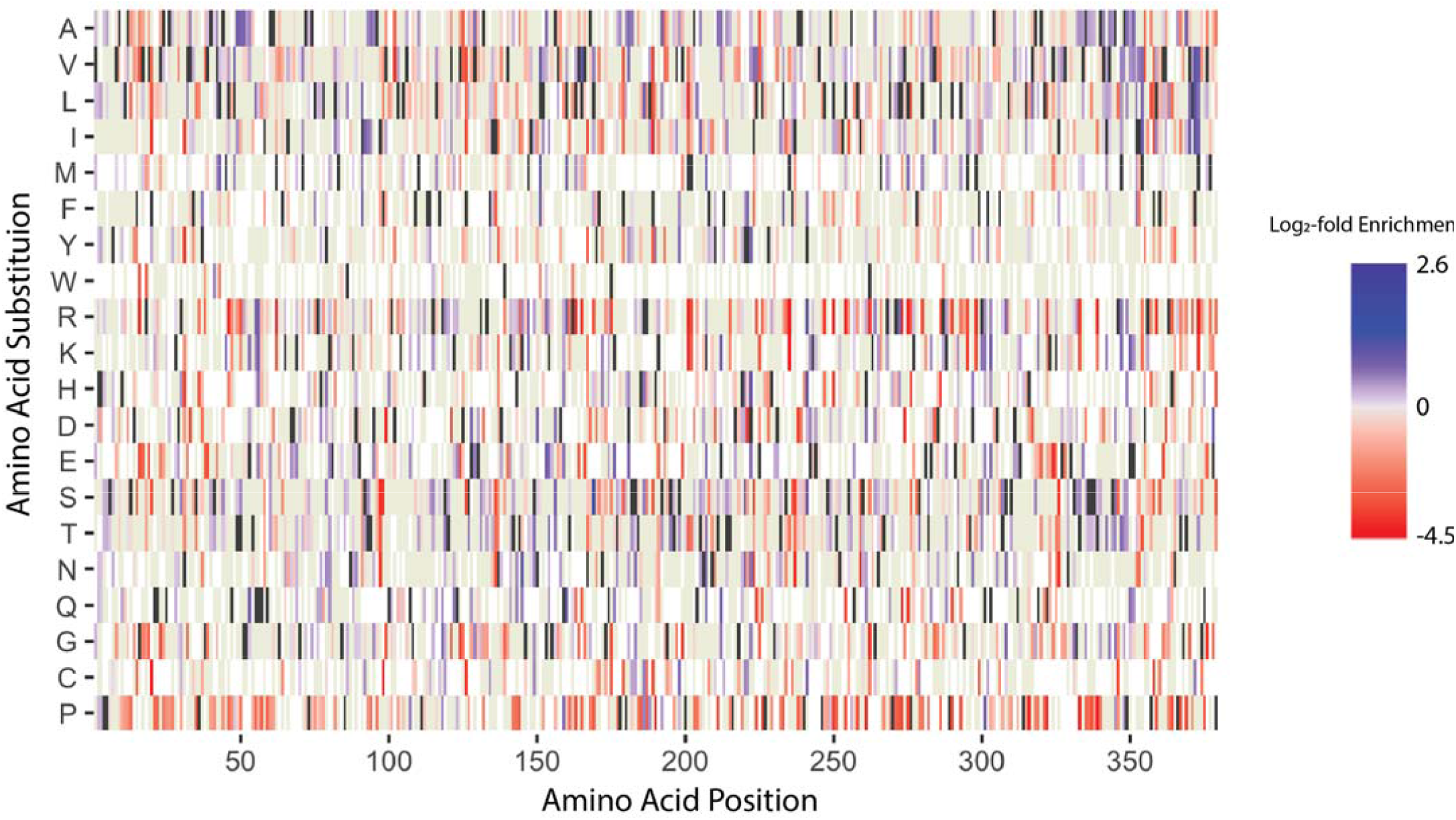
phPAI-1 mutational libraries contain missense variants that result in loss of function. Amino acid position is indicated on the x-axis, while amino acid substitutions are indicated on the y-axis. Loss of function missense variants, as well as those with a reduced capacity to inhibit uPA, are shown in red with shading as a function of their log_2_-fold score, while variants in blue retain PAI-1 inhibitory function. The intensity of shading scales with the degree of enrichment or depletion. WT amino acid residues are shown in black, while beige indicates missense mutations that were present in the mutational library but did not meet significance thresholds. White indicates amino acid substitutions that were not present in the mutational library.

### Implications for structure-function relationships

The results of our PAI-1 functional screen in the region of the RCL (residues 331-350), are illustrated in Fig. 6. The observed enrichment and depletion at the P1 and P1’ positions (residues 346 and 347) are consistent with our understanding of PAI-1 biology. The P1 position has been shown to be a key determinant of SERPIN target protease specificity (28-32), with PAI-1 inhibitory activity toward uPA requiring either a P1 Lys or the WT Arg residue (33) Consistent with these previous reports, no missense mutations were tolerated at P1 in our screen (of note, lysine at this position is absent from our library), with several substitutions significantly depleted (Fig 3 and Fig 6). Consistent with the previously reported tolerance of the P1’ position for most amino acid substitutions (33), our screen identified no loss-of-function PAI-1 variants at this position (Fig 6).

**Figure 6.**
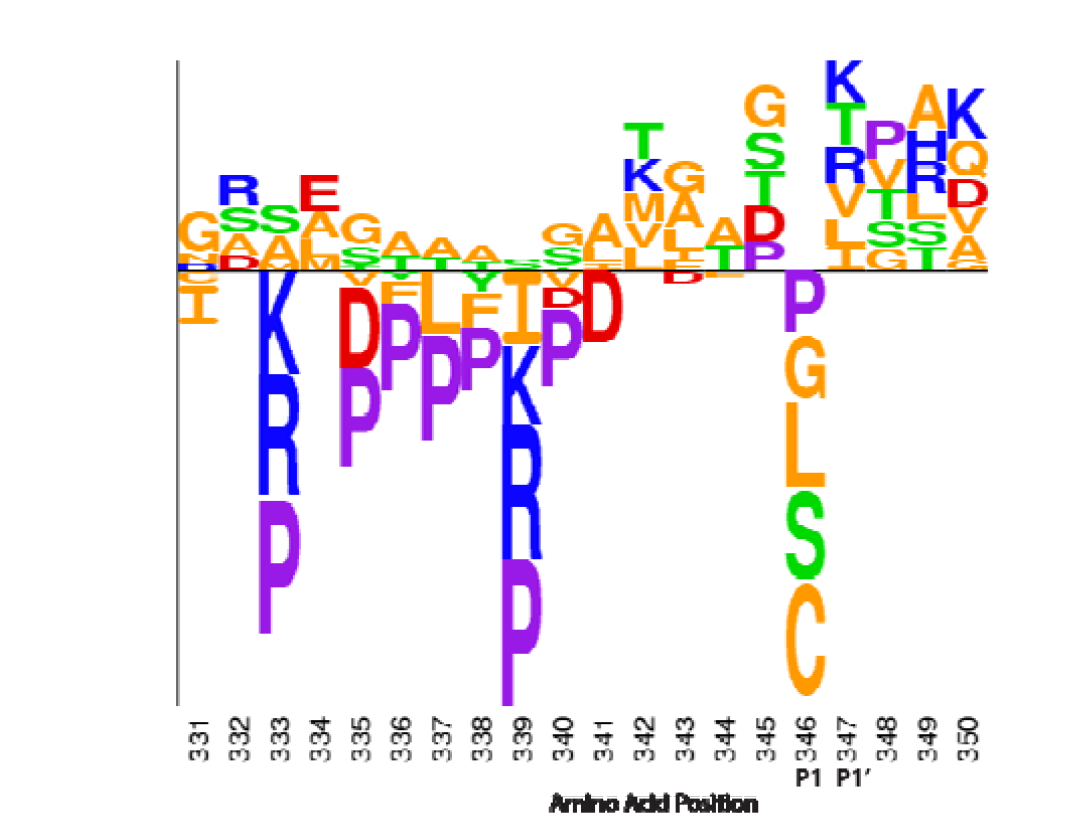
Deep mutational scanning of PAI-1 provides insight into the mutational landscape of PAI-1’s RCL. Relative log_2_-fold change scores of missense mutations in PAI-1’s RCL (amino acids 331-350) following selection by ability to inhibit uPA (31). Amino acids below the x-axis indicate depletion (log_2_-fold change < 0), while those above the x-axis indicate enrichment (log_2_-fold change > 0). Letter height corresponds to the relative log_2_-fold change. Residues are color coded by properties: acidic residues in red, basic residues in blue, polar amino acids in green, non-polar amino acids in orange, and Pro in purple (31). The P1 and P1’ positions (Met^346^ and Arg^347^ in WT) are also indicated.

At the N-terminus of the RCL, enriched or tolerated substitutions observed in our data generally consist of small aliphatic and polar amino acids. For PAI-1 to retain its inhibitory function, this region of the RCL must be able to insert into □-sheet A (9). These small amino acids allow the RCL to undergo the dramatic conformational changes that are required for this insertion. Consistent with this model, substitutions with bulky and/or charged side chains (Lys, Arg, Pro, Asp, Phe) were the most depleted residues at positions in which the RCL side chains become oriented into the core beta sheet upon insertion (34-37). In contrast, residues C-terminal to the scissile bond (P2’-P4’) are more tolerant of mutations than those at the N-terminus of the RCL, as the former region does not insert into the central β-sheet (38). Finally, the flexibility of the RCL is also important for dictating PAI-1’s inhibitory behavior, and our data are concordant with a previous proline-scanning mutagenesis screen (34). Proline residues in the RCL would also be incompatible with RCL insertion into β-sheet A, which transforms it from a largely parallel β-sheet to a more stable anti-parallel β-sheet.

### Correlation with predictive algorithms and human genome sequence variant data

A number of algorithms have been developed to predict the impact of single amino acid substitutions on protein function based on evolutionary conservation and/or amino acid type (39). We compared our high throughput screening data with predictions from two commonly used algorithms, SIFT (40) and PolyPhen-2 (41). SIFT predicts the effects of amino acid substitutions by comparison to homologous sequences, and PolyPhen-2 uses both sequence conservation and structural homology to predict the effects of amino acid substitutions on protein function.

The SIFT algorithm prediction was concordant for 745 of the 1137 (66 %) amino acid substitutions scored as “loss of function” in our screen and for 538 of the 826 (65%) scored as neutral. This level of concordance is similar to that previously reported for known deleterious human genetic mutations in other genes (40). PolyPhen-2 exhibited concordance with our data for 994 of 1137 (87%) “loss of function” substitutions, but only 454 of the 836 (54%) neutral PAI-1 amino acid substitutions (41).

Additionally, available human genomic sequence information provides support for the potential value of our data in interpreting the significance of human genetic variation identified by future clinical sequencing. The gnomAD database (42) catalogs human amino acid sequence variant information from ∼140,000 human exomic/genomic sequences, including 202 variants scored in our mutation screening analysis. Of these 202 variants, 92 were classified by our data as “loss of function”, significantly less than expected by chance (p = 2×10^−4^ SI Table 3), consistent with evolutionary selective pressure in the human population to maintain PAI-1 activity.

## DISCUSSION

Deep mutational scanning has been applied to a number of proteins to analyze function, binding interactions, cellular protein abundance, cell growth/viability, and protein stability (24,25,43-45). In the present study, we have adapted this approach to construct a detailed map for the mutational landscape of PAI-1 with respect to the gain or loss of its capacity to inhibit uPA. We anticipate that the mutational landscape of PAI-1 for other serine proteases, including its other canonical substrate, tPA, would likely demonstrate significant differences (46).

Previous PAI-1 mutational studies (47) have been restricted to limited segments of PAI-1 (33,46), or selected for a few variants with a unique functional impact, such as extended functional stability (27,48). The error prone PCR approach used here to generate the phPAI-1 mutant library offers speed and ease of application with broad coverage of a significant subset of potential single amino acid substitutions. However, variant coverage is incomplete (Fig. 3), providing significant loss of function data for only a subset of mutation space (Fig 5). Advances in molecular approaches and machine learning algorithms should facilitate a comprehensive map of the mutation landscape for PAI-1 and many other proteins (49). These data should provide a valuable resource for the interpretation of sequence variants in PAI-1 and other genes identified by the expanding clinical application of human whole genome sequencing. Future PAI-1 deep mutational screening studies should provide additional insight into PAI-1 structure-function relationships and the determinants of SERPIN target protease specificity.

## EXPERIMENTAL PROCEDURES

### Materials

Restriction enzymes, protein G magnetic beads, enterokinase, and NEBNext DNA Library Prep Master Mix Kit were purchased from New England Biolabs (Ipswitch, MA). The GeneMorph II Random Mutagenesis Kit, QuickChange II XL Site-Directed Mutagenesis Kit, and chemically and electrocompetent XL-1 Blue MRF’ *E. coli* were obtained from Agilent Technologies (Santa Clara, CA). LB broth and anti-c-myc agarose gel were purchased from Invitrogen (Waltham, MA). M13KO7 helper phage were obtained from GE Healthcare (Chicago, IL). Ampicillin, kanamycin, 3X FLAG peptide, and anti-FLAG agarose beads were purchased from Sigma-Aldrich (Saint Louis, MO). High molecular weight (HMW)-uPA was obtained from Molecular Innovations (Novi, MI). EDTA-free protease inhibitor cocktail and isopropyl-β-D-thiogalactoside (IPTG) were purchased from Roche (Basel, Switzerland). A polyclonal rabbit-anti-uPA antibody (ab2412) was purchased from Abcam (Cambridge, UK). Z-Gly-Gly-Arg-AMC is a product of Bachem (Bubendorf, Switzerland). NEXTflex multiplex adapters were purchased from Bioo Scientific (Austin, TX).

### Construction of a phage display library expressing PAI-1 fusion proteins

For display of PAI-1 on the M13 filamentous bacteriophage (phPAI-1, Fig. 1B), human SERPINE1 cDNA including a N-terminal myc tag and Gly-Gly-Gly-Ser linker was cloned between the AscI and NotI restriction sites of pAY-FE (Genbank #MW464120) (50,51). The resulting construct encodes a phage-displayed PAI-1 protein N-terminally fused to a myc tag and C-terminally fused with FLAG and E tags (Fig. 1). The PAI-1 fusion protein was randomly mutagenized using the GeneMorph II Random Mutagenesis Kit. Primers used for PCR mutagenesis (SI Table 1) maintained the AscI and NotI restriction sites for ligation of the restriction digested insert into pAY-FE. Following ligation, the library was transformed into electrocompetent XL-1 Blue MRF’ *E. coli* as per manufacturer’s instructions. The depth of the library was determined by quantifying the number of ampicillin resistant colonies. Mutation frequency was estimated by Sanger sequencing of the SERPINE1 inserts from randomly selected individual colonies (n=13).

### Phage production and purification

Phage were prepared as previously reported (52) with modifications to minimize PAI-1 conversion to the latent conformation. Briefly, *E. coli* harboring pAY-FE PAI-1 were grown in LB Broth supplemented with 2% glucose and ampicillin (100 μg/mL) at 37°C and during mid-log phase (OD_600_ 0.3-0.4) were infected with M13KO7 helper phage at a multiplicity of infection of ∼100, followed by growth for an additional 1 h at 37°C. Cells were pelleted by centrifugation (4250xg for 10 m at 4°C), resuspended in 2xYT media (16 g/L tryptone, 10 g/L yeast extract, 5 g/L NaCl) supplemented with ampicillin (100 μg/mL), kanamycin (30 μg/mL) and IPTG (0.4 mM) to induce expression of the PAI-1 fusion protein, and grown for 2 h at 37°C (50). All subsequent phage preparation steps were carried out at 4°C. Phage were precipitated with polyethylene glycol-8000 (2.5% w/v) and NaCl (0.5 M) for up to 16 h (with confirmation of no change in the number of active phPAI-1 present after purification) followed by centrifugation (20,000xg for 20 min at 4°C). The precipitated phage pellet was resuspended in 50 mM Tris containing 150 mM NaCl (pH 7.4; TBS). Phage titer was determined by transducing naïve XL-1 Blue MRF’ *E. coli* grown to mid-log phase for 1 hour at 37°C and plating on LB-agar supplemented with ampicillin (100 μg/mL) and 2% glucose.

### Selection of uPA-bound phPAI-1

Phage displaying the A3 domain of von Willebrand Factor (phVWF-A3), in which the VWF A3 domain (Ser1681-Cys1872) was PCR amplified (SI Table 1) and cloned into the pAY-FE vector between the AscI and NotI restriction sites (generating pAY-FE-VWF-A3), were used as a negative control for uPA binding. phPAI-1 were diluted 9:1 with phVWF-A3 and then incubated with uPA (1.7 nM) for 30 minutes at 37°C. Residual protease activity was inhibited by incubating the reaction mixture with 1X EDTA-free protease inhibitor cocktail for 10 min at 37°C. uPA (free and complexed) was immunoprecipitated using magnetic protein G beads (15 □L), which were previously coupled to a polyclonal anti-uPA antibody (17 nM). Beads were washed four times with TBS containing 5% BSA (1 mL), resuspended in Tris (20 mM) pH 8.0 containing 50 mM NaCl, 2 mM CaCl_2_, and 5% BSA, and eluted by digestion with enterokinase (16 U) for 16 hours at 4°C. The eluted phage pool was used to infect naive XL-1 Blue MRF’ *E. coli*. Eluted phage titers were quantified by transduction of XL-1 Blue MRF’ cells as described above. To determine the composition of the phage pools before and after selection, single colonies of ampicillin resistant bacteria were selected, and their DNA amplified by PCR using primers annealing outside the insertion site, to a region common to both pAY-FE:PAI-1 and pAY-FE:VWF-A3 (SI Table 1) with three replicates of n = 24 colonies in each.

### High-throughput sequencing (HTS)

Twelve overlapping amplicons (150 bp) were PCR amplified from pAY-FE PAI-1 (SI Table 1) with overlapping regions only analyzed on one amplicon (SI Table 2). PCR amplicon products were gel purified, pooled (100 ng DNA divided equally between 12 amplicons), dA-tailed (NEBNext Ultra End Repair/dA-tail), and ligated to NextFlex barcodes with NEBNext Ultra Ligation. Ligated Products were purified with AmPure beads according to the manufacturer’s directions. HTS was performed as previously described (25) using the Illumina HiSeq2500 or HiSeqX platforms (Illumina, San Diego, CA) at the University of Michigan DNA Sequencing Core (Ann Arbor, MI) or MedGenome, Inc (Foster City, CA). HTS data were analyzed using DESeq2 (26) analyzing mutations at each position independent of other mutations within a given amplicon.

### Comparison of variant selection results to publicly available datasets

Results of the DESeq2 analysis of input versus selected were compared to the output from the Sorting Intolerant from Tolerant (SIFT) algorithm that predicts the effect of an amino acid substitution on protein function by multiple sequence alignments of related proteins (PAI-1 from *S. scrofa, B. taurus, M. vison, R. norvegicus*, and *M. musculus;* glia-derived nexin from *H. sapiens, M. musculus, and R. norvegicus;* neuroserpin from *H. sapiens, G. gallus, R. norvegicus, and M. musculus*) (53). To compare SIFT results to our data, tolerated mutations were defined as those that were able to inhibit uPA (log_2_-fold change > 0) at 0 h, and noninhibitors (log_2_-fold < 0) at 0 h were defined as not tolerated. Finally, a χ^2^ test (SI Table 3) was used to determine if the variants identified as loss-of-function by our high-throughput screen were significantly underrepresented in the gnomAD database (42) by comparing the expected frequency of variants identified in our screen that were present in gnomAD versus those that were not present.

## DATA AVAILABILITY

Sequencing data will be publically available following publication and made available at the reviewers’ request.

## SUPPORTING INFORMATION

This article contains supporting information.

## FUNDING AND ADDITIONAL INFORMATION

This work was supported by the National Institutes of Health grants R35-HL135793T (DG) and R01-HL055374 (DAL), T32-GM007863 (ZMH), and T32-HL125242 (ZMH). LMH was supported by a National Hemophilia Foundation Judith Graham Pool Postdoctoral Research Fellowship. AY is supported by and American Society of Hematology Faculty Scholar Award, National Hemophilia Foundation Innovative Investigator Research Award, and Mary R. Gibson Foundation. CAK is supported by an Early Career Award from Hamilton Health Sciences. DG is a Howard Hughes Medical Institute investigator. The content is solely the responsibility of the authors and does not necessarily represent the official views of the National Institutes of Health.

## SUPPORTING INFORMATION

**SI Table 1.**
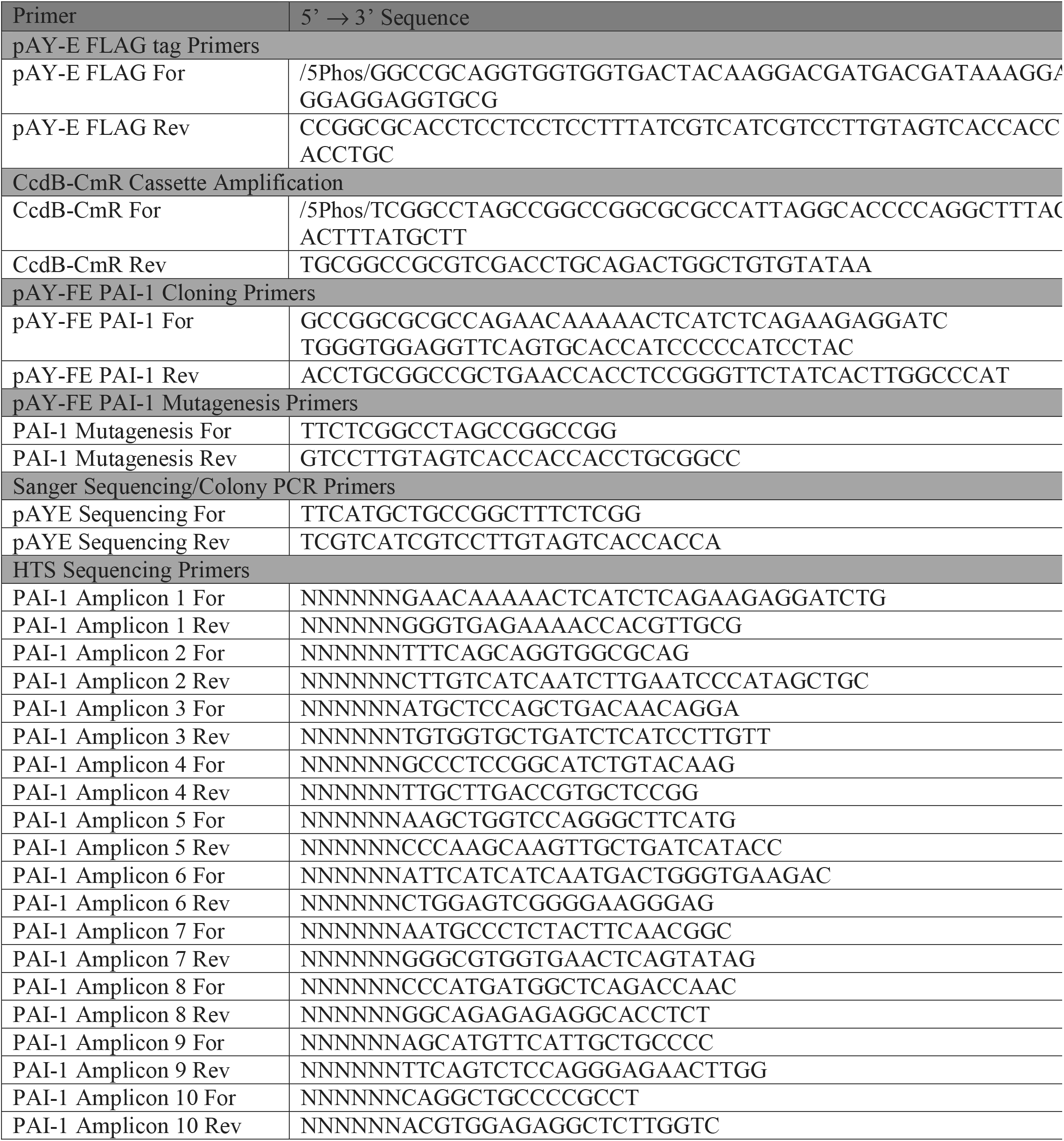
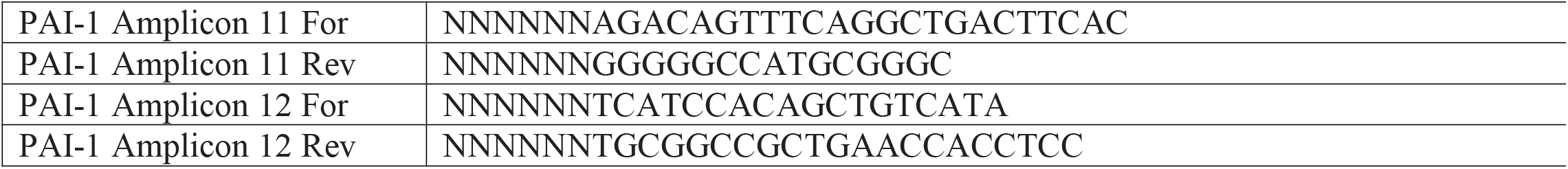
Primer Sequences

**SI Table 2.**
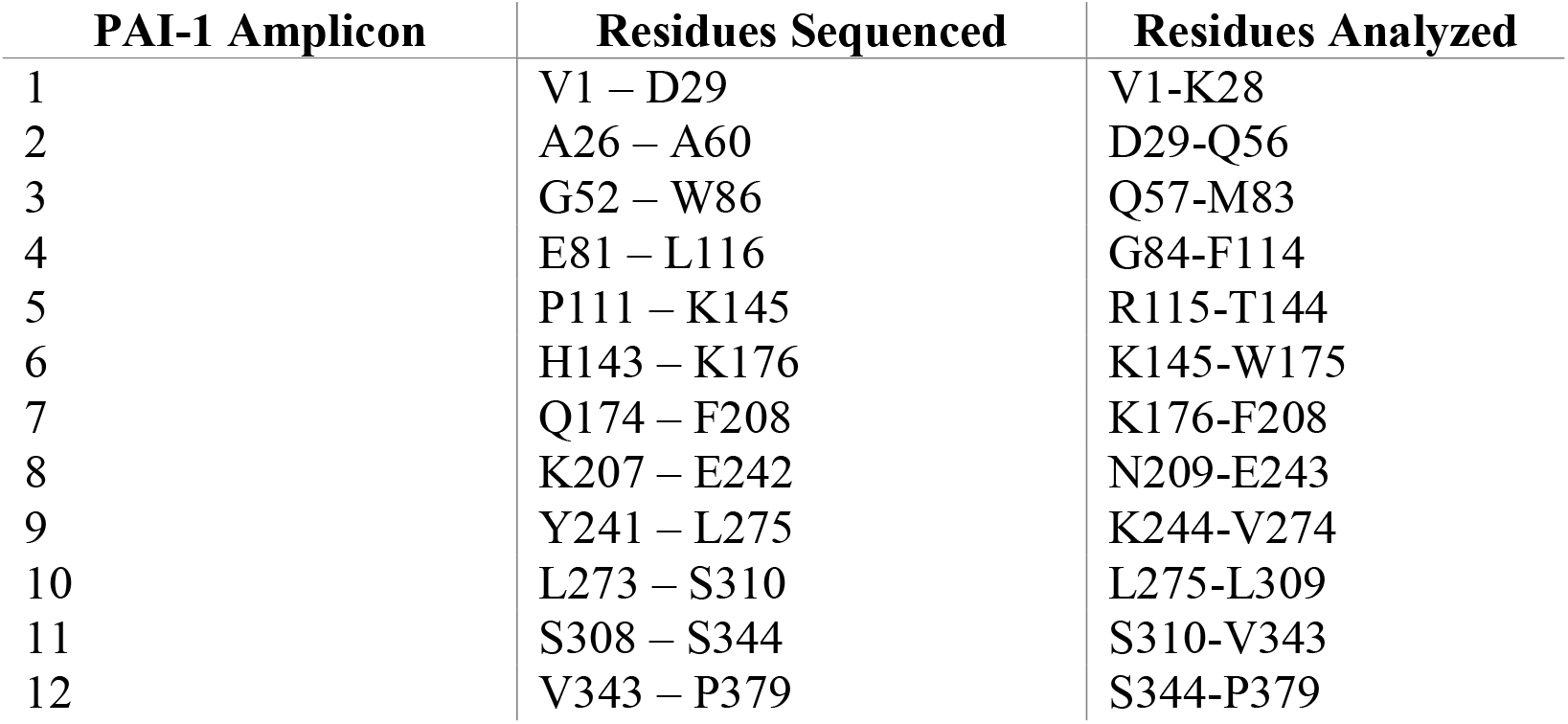
PAI-1 amino acid residues spanned by HTS amplicons

**SI Table 3.**
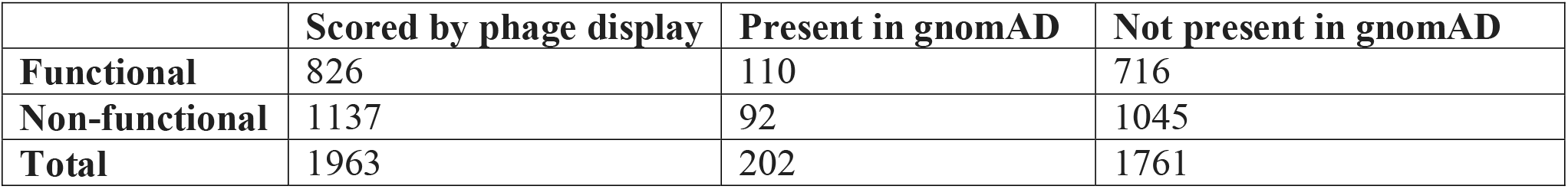
Observed frequencies for χ^2^ test comparing experimental results to gnomAD database

